# Reorganization of Pancreas Circadian Transcriptome with Aging

**DOI:** 10.1101/2023.05.17.541196

**Authors:** Deepak Sharma, Caitlin R. Wessel, Mahboobeh Mahdavinia, Fabian Preuss, Faraz Bishehsari

## Abstract

The evolutionarily conserved circadian system allows organisms to synchronize internal processes with 24-h cycling environmental timing cues, ensuring optimal adaptation. Like other organs, the pancreas function is under circadian control. Recent evidence suggests that aging by itself is associated with altered circadian homeostasis in different tissues which could affect the organ’s resiliency to aging-related pathologies. Pancreas pathologies of either endocrine or exocrine components are age-related. Whether pancreas circadian transcriptome output is affected by age is still unknown. To address this, here we profiled the impact of age on the pancreatic transcriptome over a full circadian cycle and elucidated a circadian transcriptome reorganization of pancreas by aging. Our study highlights gain of rhythms in the extrinsic cellular pathways in the aged pancreas and extends a potential role to fibroblast-associated mechanisms.

## Introduction

Optimal biological functions of an organism are highly dependent on its ability to adapt to environmental pressures. Because of the rotation of the Earth around its own axis and the sun, life evolved under multiple rhythmic environmental regimes, including both seasonal and daily changes in light exposure^1^. As a result, nearly every species on Earth has developed a biological timekeeping system that can anticipate these changes and organize physiology and behavior in a way that is advantageous for the organism. Biological rhythms of approximately 24 h, called circadian rhythms, are highly conserved among mammalian species and provide a temporal order for behavioral and physiological processes^2^.

Almost all physiological processes in mammalian organs are under circadian regulation^3,4^. The circadian rhythm apparatus consists of a central master clock located in the suprachiasmatic nuclei (SCN) within the hypothalamus, which synchronizes peripheral oscillators located in various tissues such as the liver, pancreas, adipose tissue, and skeletal muscle. The SCN is entrained by light^5,6^, which allows synchronization with the external environment i.e., the 24-hour light-dark cycle governed by the Earth’s rotation. This directs the central and peripheral clocks to adapt to changes in light, optimizing physiological processes to these daily cycles. Through several regulatory mechanisms (e.g., endocrine, neurological, thermal), the SCN coordinates responses with the peripheral clocks, which have their own phases, to ensure synchronized daily rhythms are maintained^7^. In addition, peripheral rhythms can also be modulated, for example, by nutrient sensing (i.e., from food intake), hormonal cues and temperature^8^. Although the SCN acts as the master pacemaker in the human body, it is evident that circadian oscillations are observable in almost every cell of the body and these rhythms may persist in isolation from the SCN^8^.

Studies have shown that pancreas function is strongly under circadian control and disruption of circadian rhythms exacerbates metabolic diseases that include Type II diabetes mellitus, obesity and metabolic syndrome in both animal models and humans^9-12^. In addition, circadian functions are known to decline over the lifespan in several organs^13^. For example, age-related changes in the amplitude of sleep/wake activity, body temperature, metabolic functions, and hormone release is well documented^14^. Aging is also connected with the inability to adjust to a new light/dark cycle, and increased mortality following repeated jet lag^15-16^. A few recent age-related studies showed that the core clock in aged tissues remains largely intact, while the transcriptional output is significantly altered^17^. Despite these studies, a systematic and mechanistic study of the age-dependent temporal gene expression profile of pancreas is largely lacking. Given the strong connection between age and the resiliency of pancreas to disease (both exocrine and endocrine pathologies), knowledge of differential pancreatic circadian transcriptome could lead to deeper understanding of its function and unearth novel targets for therapeutic interventions against age-related pancreas pathologies such as metabolic disorders and cancers.

Here we carried out a 24-h circadian transcriptomic analysis of pancreas from male mice at young and old ages. We define a comprehensive circadian transcriptome landscape and identify biological pathways that are reflective of aging pancreas. Additionally, analysis of pancreatic microenvironment revealed novel mechanistic insights into the fibroblast mediated regulation of rhythmicity in aged pancreas. We suggest that circadian transcriptome in aging pancreas re-organizes in response to age-specific signals from the cellular microenvironment, primarily modulated by fibroblasts.

## Results

### Changes in steady state gene expression in pancreas with aging

To quantitate the effects of aging on the molecular landscape of pancreas, we profiled the gene expression of the mice pancreas using RNA-seq (depth: >40 Mi reads). Two age-groups of male C57/BL6J mice were entered to our protocol (see method). Briefly, mice at 4 months or 24 months of age were singly housed and entrained under 12Light:12Dark conditions for 1 month. After the end of light phase of the last day of entrainment the lights remained off and the pancreatic tissues were collected in 4 hours intervals (6 time points, n = 2 mice per time point) starting at ZT0 in constant dark, at 5 months or 25 months of age. The transcriptome profiles for each time point were highly similar between age groups (R^2^ > 0.95; Figure 1A). However, visualization over 24-h revealed clustering of young and old group separately, indicative of temporal molecular profiles (Figure 1B). Aged compared to young pancreas showed an upregulation in 133 genes and downregulation of 251 genes (p < 0.05; fold change ≥ 1.5) (Figure 1C and Supplementary Figure 1). Upregulated genes were enriched (FDR; p < 0.05) in pathways related to membrane transport and metabolism, consistent with the changes reported in the process of aging^18^ as well as the extracellular matrix (ECM) organization, while downregulated genes were enriched in pathways connected with the immune regulation and signaling (Rho GTPase, NF-κB) (Figure 1D).

**Figure 1:**
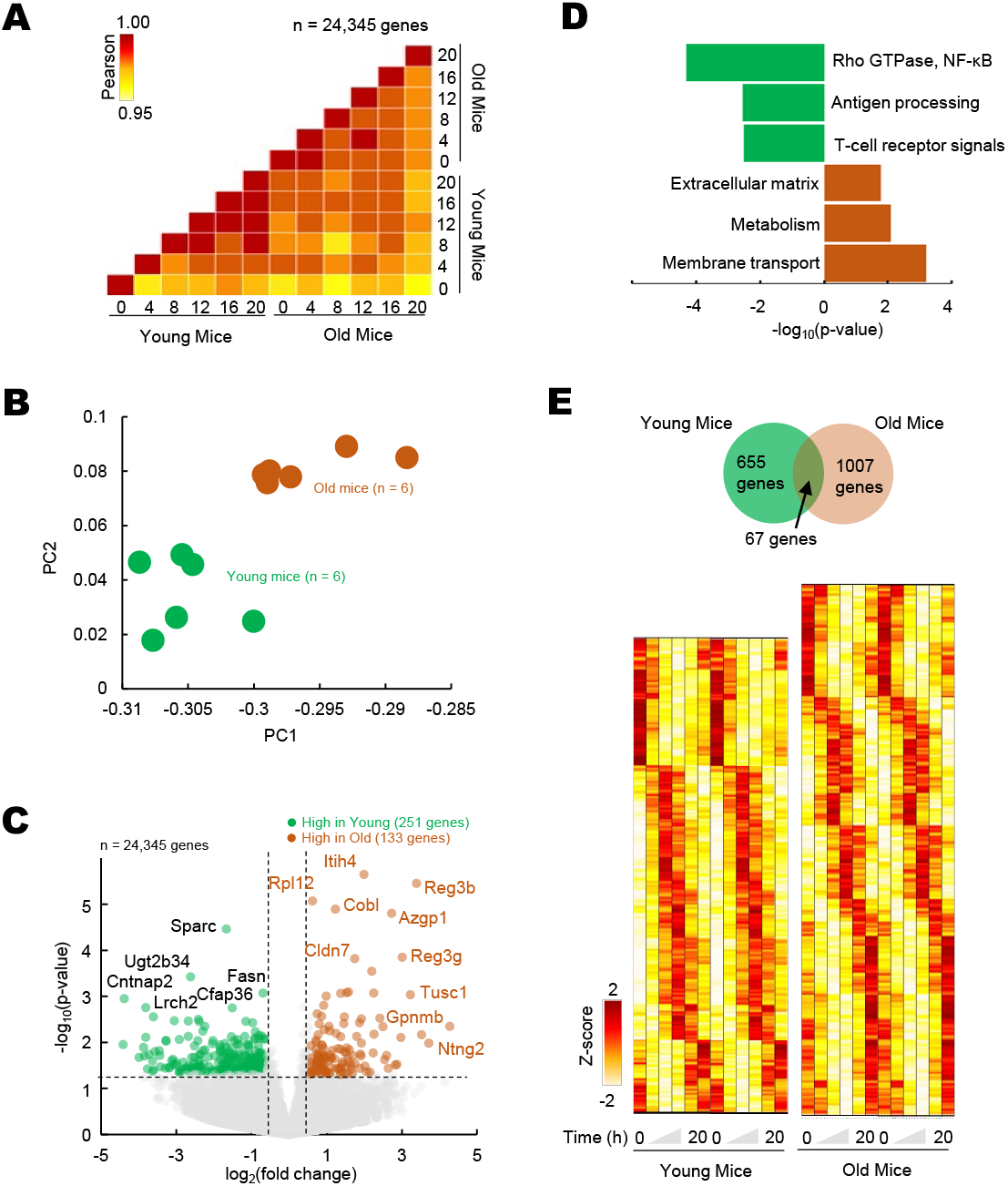
Differential and temporal gene expression signatures in aged pancreas. (A) Pairwise correlation of normalized gene expression (transcripts per million; TPM) for young and old RNA-Seq samples (n=12). Colors indicate the Pearson correlation. (B) Principal Component Analysis (PCA) of temporal gene expression from young and old mice samples (n = 12). (C) Top: Volcano plot showing differential gene expression profile with significantly (p < 0.05, fold change ≥ 1.5) upregulated (n = 133 genes) and downregulated (n = 251) genes in old mice when compared to young. (D) Enriched molecular pathways corresponding to up and downregulated genes. (E) Top: Venn diagram of rhythmic genes in young and old mice. Bottom: Heatmap of z-scored rhythmically expressed genes in young and old mice. Heatmaps are double plotted for time.

### Re-organization of circadian cycling of gene expression in aged pancreas

To delineate the differential circadian transcriptome of pancreas by age, we defined cycling genes of transcripts with 24-h cosine oscillations (FDR; p < 0.05, Meta2D)^19^. We identified only 67 overlapping rhythmic genes, indicating that pancreas circadian transcriptome is age specific (Figure 1E). Notable rhythmic genes in both subgroups were the core clock genes, *Arntl, Per2, Tef*. On the other hand, genes involved in ECM reorganization, *Col1a1* and *Cdh11* were rhythmic only in old pancreas, while genes involved in mRNA metabolism were rhythmic in younger pancreas (e.g., *Cpsf3*, Figure 2A). Pathway enrichment analysis of rhythmic genes showed an enrichment in cell intrinsic processes (RNA metabolism, DNA replication) in young pancreas, while in the old group, the cell extrinsic cellular processes (ECM re-organization, collagen formation) were rhythmic (Figure 2B). Temporal pathway enrichment showed distinct processes enriched at different time points in young and old pancreas (Figure 2C), indicative of unique age specific circadian profiles. We then asked if this distinct circadian transcriptome could be due to age-dependent central circadian cues or molecular clock dysfunction. To assess the molecular clock, we first analyzed the hypothalamus, the location for the central circadian clock. The clock correlation distance (CCD)^20^ of the temporal relative expression of 12 core clock genes revealed highly similar core clock profiles in both age groups (Supplementary Figure 2). An identical pattern also existed in the pancreas where clock CCD between young and old mice were similar (Figure 2D). These observations indicate that the core circadian clock progression is overall conserved, while extrinsic cellular processes gain rhythmicity in aged pancreas.

**Figure 2:**
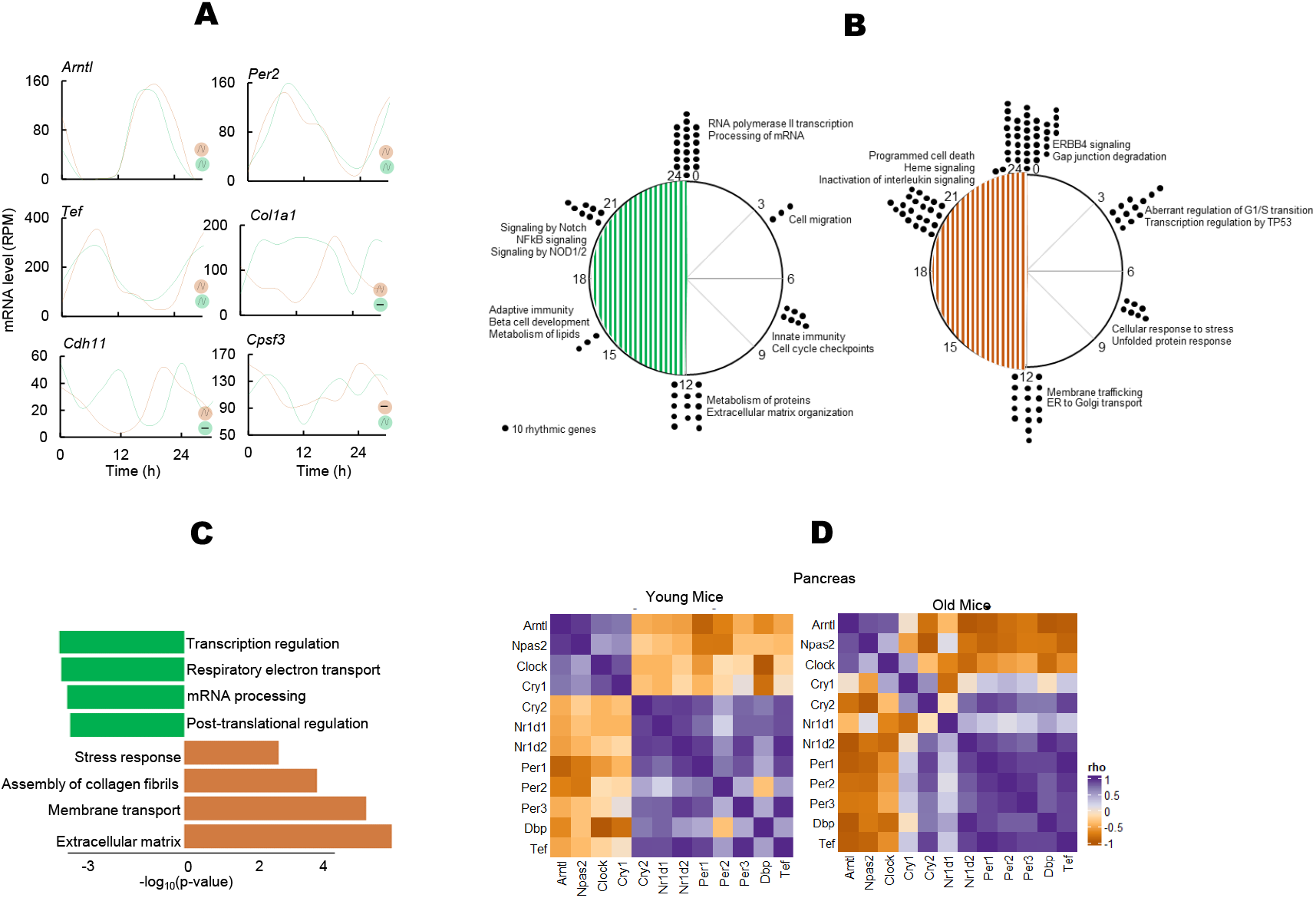
Circadian transcriptome of the aged pancreas. (A) Examples of circadian profiles of genes that are rhythmic in both young and old mice (Arntl, Per2, Tef), only in old mice (Col1a1, Cdh11), and only in young mice (Cpsf3). (B) Enriched molecular pathways corresponding to rhythmic genes in young and old mice. (C) Peak time maps of all oscillating genes from young and old mice. Each dot represents the peak time of 10 significantly rhythmic genes. Enriched pathways corresponding to each time point are also shown. (D) Clock correlation distance (CCD) of 12 core clock genes from young and old mice pancreas.

### Fibroblast mediated age-modified cellular microenvironment is a hallmark of differentially rhythmic, aged pancreas

Extrinsic pathways are known to be modulated by the cellular microenvironment of interacting soluble factors and signals in the ECM consisting of stromal and immune cells^21^. We reasoned that the observed rhythmicity and upregulation in extrinsic pathways in aged pancreas could be a result of an age-modified tissue microenvironment. We imputed the gene signature based fractional abundances of 64 immune and stromal cell types from the old and young pancreas (Supplementary Figure 3A). Fibroblasts was estimated to be the most significantly altered cell type by far with markedly higher abundance in the aged pancreas (Figure 3A, p < 10^−8^). Fluorescence Activated Cell Sorting (FACS) of the remaining pancreas of the studied mice conclusively showed significantly more activated fibroblasts in old mice (Figure 3A, p < 10^−9^ and Supplementary Figure 3B). This data suggests that fibroblasts are heavily involved in the aging of pancreas, which is consistent with the known role of fibroblasts in aging^22^ and points towards a fibroblast associated reorganization of pancreatic microenvironment. Using datasets of 178 human pancreas samples from Genotype-Tissue Expression (GTEx) project, we found a significant increase in fibroblast fractions in aged human pancreas (Figure 3B), indicating that this pattern is conserved across the species.

**Figure 3:**
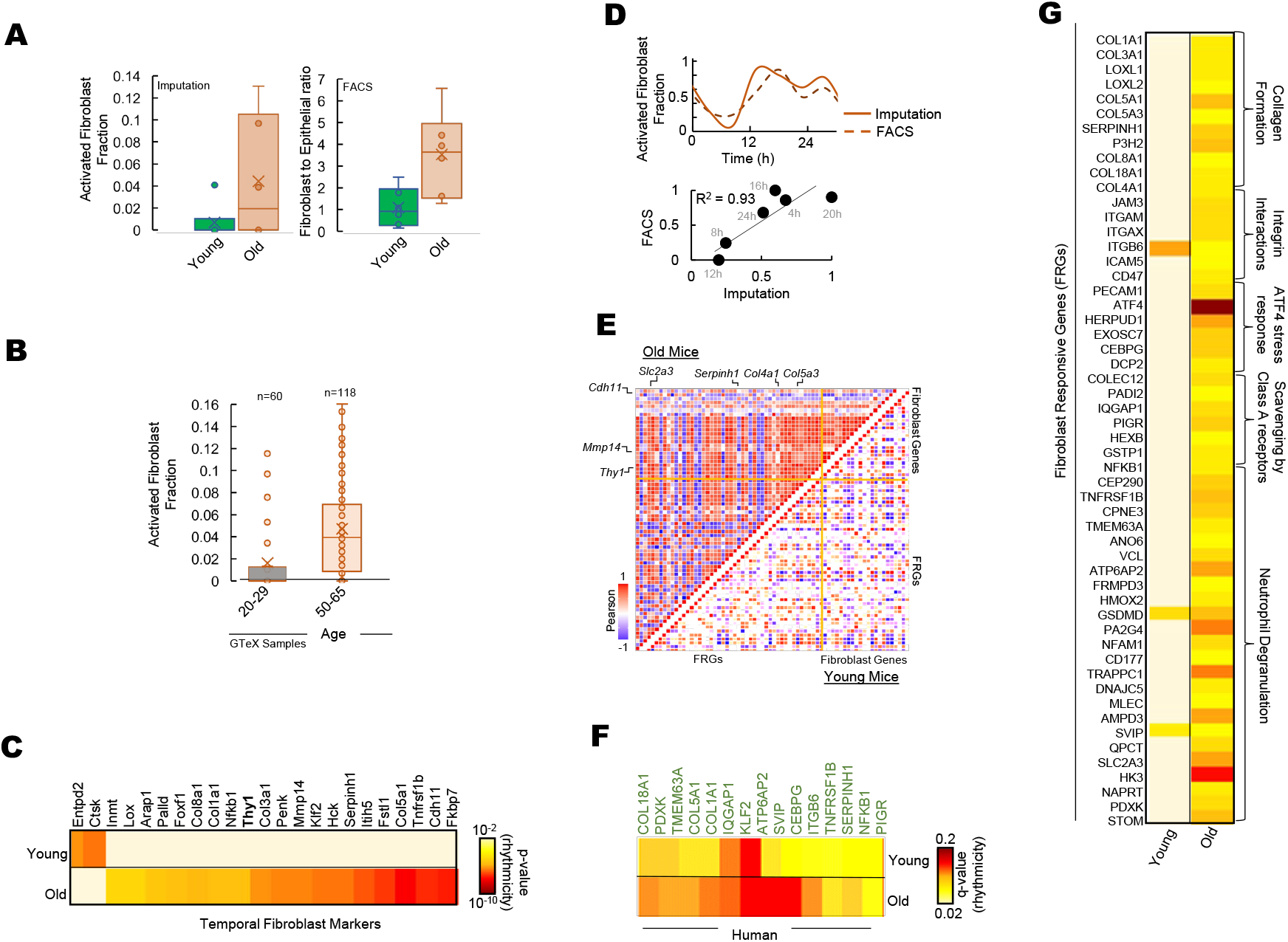
Fibroblasts mediate differential rhythmicity in aged pancreas. (A) Bar plots for (left) activated fibroblast fraction obtained by imputation and (right) Fibroblast to epithelial ratio obtained by Fluorescence Activated Cell Sorting (FACS) analysis for young and old mice pancreas. (B) Bar plots for activated fibroblast fraction obtained by imputation for young and old human pancreas from 178 GTeX datasets. (C) Heatmap of significantly rhythmic genes indicative of fibroblast markers in young and old mice. Activated fibroblast marker Thy1/CD90 is shown in bold. Heatmap represents the p-value of rhythmicity. (D) Top: Comparison of temporal activated fibroblast fractions obtained by two independent methods: imputation and FACS. Bottom: Similarity between the estimated fibroblast fractions between imputations and FACS methods. (E) Pairwise correlation plot for fibroblast markers (panel C) and FRGs (panel G) in young (lower diagonal) and old (upper diagonal) mice. Colors represent the Pearson correlation coefficient. (F) Heatmap of significantly rhythmic fibroblast responsive genes (FRGs) in young and old mice from time-stamped human datasets. Heatmap represents the p-value of rhythmicity. (G) Heatmap of significantly rhythmic fibroblast responsive genes (FRGs) in young and old mice. Heatmap represents the p-value of rhythmicity.

### Fibroblast gene activity is rhythmic in aged pancreas

We hypothesized that in aged pancreas, fibroblasts in the microenvironment may be involved in re-organization of the rhythmic processes observed in Figure 2B. We assessed the temporal expression of genes reflective of activated fibroblasts^32^. Interestingly, signatures for activated fibroblasts significantly encompassed rhythmic genes in aged pancreas (p < 10^−5^, Figure 3C), with majority of hub proteins rhythmic including the activated fibroblast marker *CD90/Thy1* (Supplementary Figure 3B and 4). No enrichment was observed in young pancreas. While there was a strong correlation between temporal fibroblast fractional estimates and FACS-based activity markers, they showed rhythmicity only in the old mice with similar 24-h cosine oscillations (Figure 3D).

### Responsive genes to fibroblasts gain rhythm in the aged pancreas

Next, we speculated that genes that are responsive to fibroblast activity should show a similar, distinct pattern in the old pancreas. Using a panel of genes responsive to activated fibroblast signals^32^ (Fibroblast Responsive Genes; FRGs, see methods), we found majority of FRGs depicted rhythmicity in aged pancreas (Figure 3G and Supplementary Figure 4). The rhythmic FRG enriched pathways were related to ECM re-organization, collagen formation, and integrin signaling. In addition, pairwise comparison of temporal expression profiles of fibroblast gene signatures and FRGs revealed a striking correlation (Figure 3E) suggesting rhythmic gene activity of FRGs mirrored fibroblast gene activity in aged pancreas. To test if a similar pattern exists in human pancreas, we assessed the rhythmicity of FRGs in time-stamped human pancreas datasets (from Ref 23). A significant number of FRGs were rhythmic in old but not in young human pancreas (Figure 3F). Overall, our study identified previously unknown circadian transcriptome reorganization of pancreas by aging, characterized by a gain of rhythm in the extrinsic cellular pathways linked to fibroblast-associated mechanisms.

## Discussion

Despite evidence of strong circadian regulation of pancreatic functions as well as impact of aging on the circadian functions across several organs, the age-dependent changes in temporal gene expression in pancreas remained unclear^9-12, 14-16^. To investigate this, we profiled transcriptome-wide gene expression of aging pancreas in mice and assessed the processes and mechanisms that gain or lose rhythmicity with age.

Our data showed that despite similar differential expression levels of steady state RNA (>98%), marked age specific differences exist in the circadian transcriptome of young and old pancreas (1,662 genes rhythmic), with extrinsic cellular processes gaining rhythm in the aged pancreas. These extrinsic pathways respond to maintain the architecture of the extracellular matrix (ECM), thus creating a three-dimensional network of macromolecules that provide structural and biomechanical support to tissue^21^. Observed rhythmicity in ECM related processes specific to the aged pancreas indicates an age-specific re-organization of the transcriptome to ensure a temporal ECM integrity that could be pivotal towards pancreatic homeostasis and resilience in old age.

Interestingly, we found that the core circadian clock structure stayed largely conserved in aged pancreas, despite gains in rhythmicity of extrinsic cellular processes. This alludes to a local re-programming of the circadian transcriptome with aging in pancreas probably due to aging related stress response in the tissue^21^. Further investigation into the-cause-and-effect connections between the changes in the ECM and rhythmicity of extrinsic cellular pathways could shed important insights into the mechanism of aging and could help identify novel targets to maintain healthy pancreas activity.

As a potential candidate that is involved in the ECM re-programming, we looked at the cellular microenvironment encompassing immune and stromal cells (despite functional relevance to the endocrine function of pancreas, islet cells could not be investigated as they make up only 2% of the pancreatic mass). Activated fibroblasts were found to be the most abundant as well as the most rhythmic in aged pancreas, with direct gene-to-gene associations of rhythmicity to the extrinsic rhythmic pathways. This previously unknown temporal re-organization of pancreatic cellular microenvironment, centered on fibroblasts markers points to a fibroblast-centered circadian re-organization of the pancreas with aging.

### Limitations of the study

Our work comes with some caveats that should be acknowledged. We collected gene expression data to assess rhythmicity over 1 cycle of 24h, at every 4h. This precluded repeated observation of peak-trough rhythmicity and could underpower our ability to detect more rhythmic genes. However, all the time points were collected in duplicates and mice were kept under constant darkness throughout the collection day to minimize the effect of light-induced changes.

In addition, we looked at any changes in rhythmicity profile across age groups in the form of pattern as reflected by the enrichment of biological pathways, and not an isolated single gene. This ensures that minor fluctuations in expression counts at a given time point for a given gene would not artificially alter the inference of rhythmicity. Lastly, further investigation into the connection between different cell populations of the cellular microenvironment and rhythmicity with aging is needed to obtain a comprehensive picture of the detailed mechanism by which ECM responsive processes gains rhythmicity in aged pancreas.

## Acknowledgements

We would like to thank Darbaz Adnan, Dan Leary and Breanna Palmen for their technical and analytical help.

## Author contributions

DS: data analysis, investigation, methodology, visualization, data interpretation, manuscript drafting.

CW: investigation, visualization.

MM: methodology, data interpretation.

FP: investigation, methodology, project administration.

FB: conceptualization, funding acquisition, investigation, project methodology & administration, supervision, data interpretation, writing.

## Declaration of interests

The authors have no conflicts of interest to declare.

## Grant support

FB is supported by NIH CA279487 and CA277110; Swim Across America Foundation; DS is partly supported by Tides Foundation

## Figure Legends

**Supplementary Figure 1:**
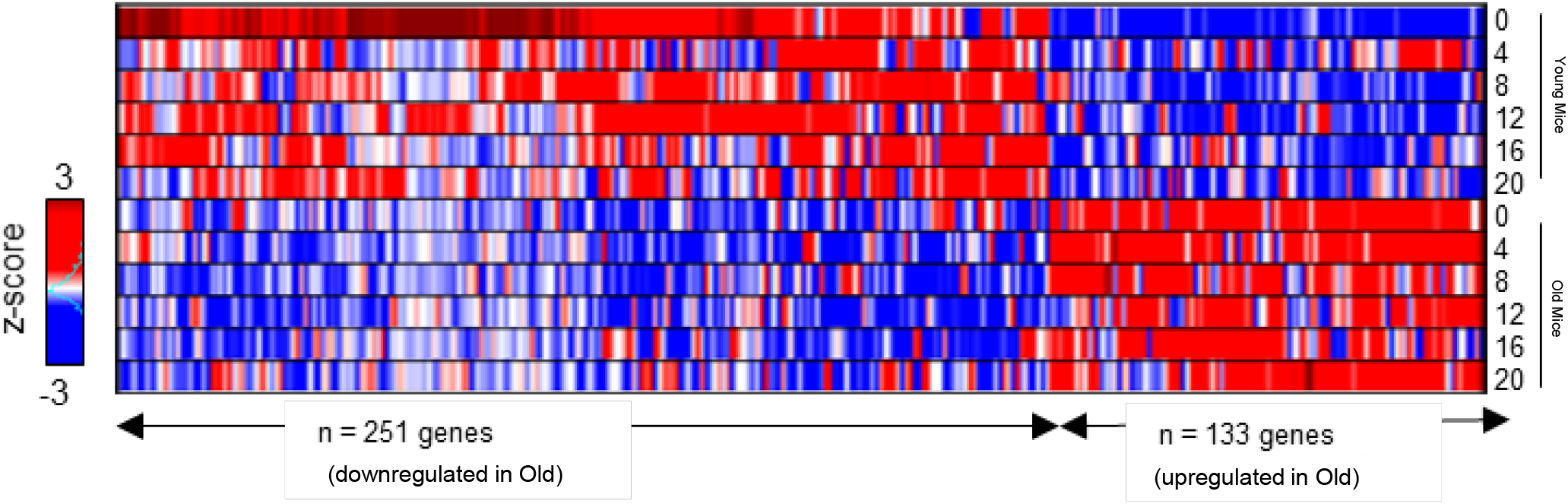
Differentially expressed genes in mice pancreas. Heatmap of z-scored expression profiles for young and old RNA-Seq samples (n=12). Colors indicate relative (inter-sample) expression profiles for significantly up and down regulated genes (from panel 1A). Red: high TPM; Blue: Low TPM; White: Similar TPM. TPM = Transcripts per Million.

**Supplementary Figure 2:**
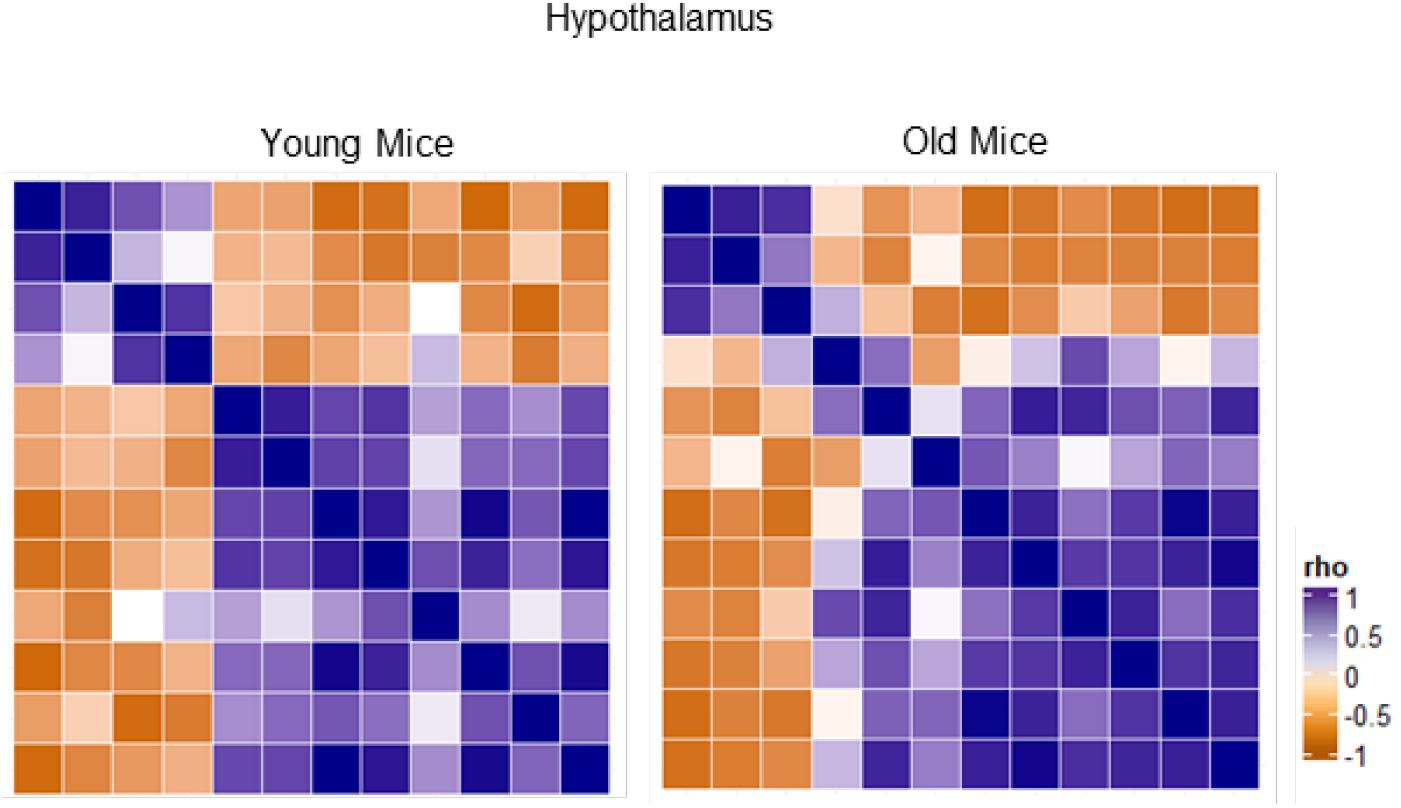
Conserved central clock with aging. Clock correlation distance (CCD) of 12 core clock genes from young and older mice hypothalamus.

**Supplementary Figure 3:**
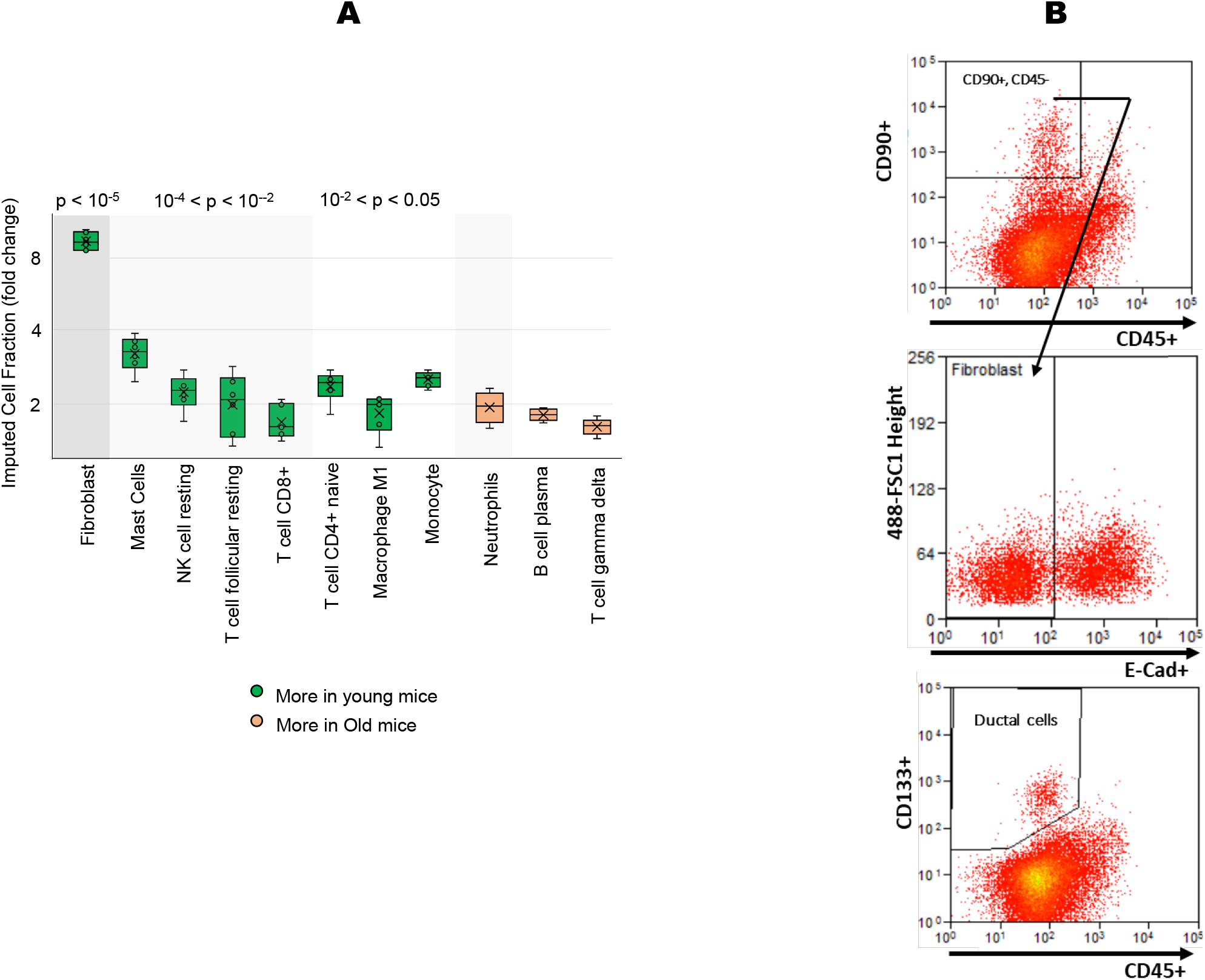
Imputation of immune cell fractions in young and old pancreas.(A) Bar plots of the fold change for activated immune and stromal cell fractions obtained by imputation for young and old mice pancreas. p-value significances are color coded in grey. Cell types not significant are not shown. Green: More cells in young compared to old mice. Brown: More cells in old compared to young mice. (B) Representative flow cytometry workflow shows an example of gating scheme for activated fibroblasts in the pancreas tissue CD90+/CD45-/E-cadherin- (upper two panels) which was then normalized to the number of ductal epithelial cells; non-hematopoietic cells (CD45-) enriched in ductal (CD133+) markers (lowest panel) in each sample.

**Supplementary Figure 4:**
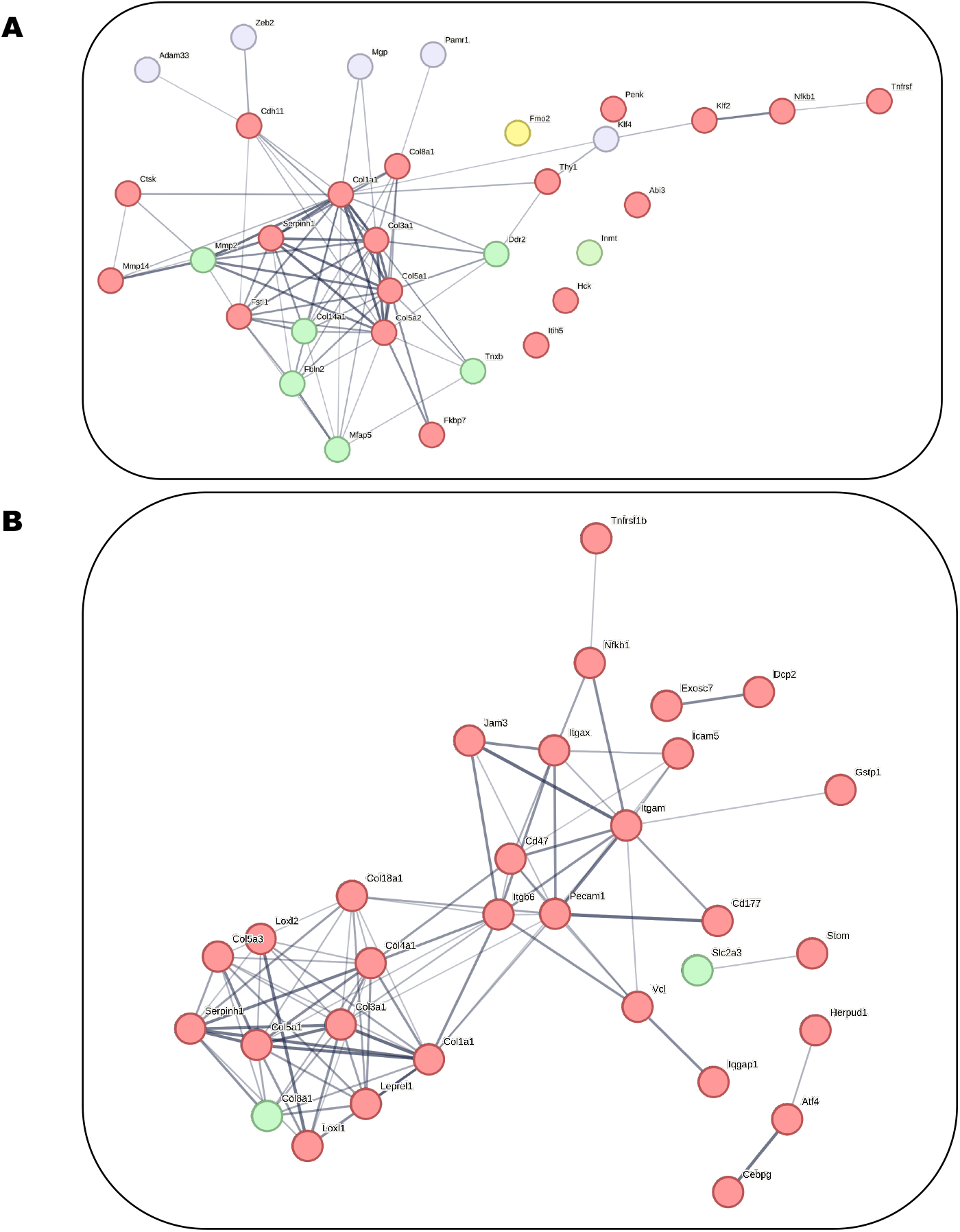
Fibroblast activity is rhythmic in aged pancreas. (A) Hub proteins corresponding to genes that are markers for activated fibroblasts. Color: Red: Rhythmic in old pancreas. Green: Rhythmic in young pancreas. Grey: No change in rhythmicity. Yellow: Rhythmic in both young and old pancreas. Hub proteins were rhythmic in old pancreas at p< 0.0001. No significance was found for young pancreas.(B) Hub proteins corresponding to genes that are responsive to fibroblast signals. The color code is the same as panel (A). Hub proteins were rhythmic in old pancreas at p< 0.0001.

## Methods

### Animal care and use

C57BL/6J mice, born and raised at the University of Wisconsin-Parkside animal facility. The parental generation was obtained from the Jackson laboratory and the described procedures approved by the institutional animal care and use committee of the University of Wisconsin-Parkside. Breeding and housing colonies were housed under standard 12 hours of light followed by 12 hours of darkness (12L:12D) in light and temperature-controlled chambers and had continuous access to regular chow (ad-libitum) and fresh water. Mice were entered to our protocol at 4 months or 24 months of age. Both groups were singly housed and entrained for 4 weeks in LD chambers. Mice were sacrificed at 5 months or 25 months of age. Animal tissues following euthanasia were collected in 4-hour intervals under dark to allow for circadian analysis. Collected samples were partly persevered in RNA later for RNA processing, while a portion was used for flow cytometry as described below.

### Sample Preparation of Total RNA Sequencing

RNA was isolated from the pancreas using TRIzol extraction procedure. Briefly, frozen tissues were lightly ground in a mortar and pestle constantly submerged in liquid nitrogen. Frozen tissue between 10-100 mg was placed into RNase-free tubes containing TRIzol and RNase-free stainless-steel beads (NextAdvance, USA). Tissues were homogenized using a bullet blender at 4°C. Chloroform was used to separate the phases, and the RNA-containing aqueous phase was subjected to a column purification kit using RNeasy Mini RNA extraction kit (Qiagen, USA). RNA was DNase treated, and purified RNA was checked for integrity using an Agilent Bioanalyzer, all samples had an RNA integrity number above 8.0. RNAseq libraries were prepared using Illumina mRNA Prep kit (Illumina, USA). Libraries were pooled to equal molarity and sequenced on an Illumina NovaSeq (2 × 100bp) to achieve a minimum of 40M reads per sample. FastQ files were downloaded to the Rush University computing cluster for further data processing.

### Flow cytometry

Pancreatic tissue was homogenized, and cells were suspended in DMEM and 10% fetal bovine serum (FBS). Supernatant was centrifuged at 500g for 5min, and the pellet was resuspended in a tissue digestion buffer (collagenase IV, dispase, DNase I, and DMEM). After arresting tissue digestion, the solution underwent centrifugation at 500g for 5min and the pellet was resuspended in DMEM, DNase 1, and 10% FBS. Total live cell retrieval varied based on timepoints, however ranged from 1.2-6.5 x 106 cells. Dead cells were excluded using the Live/Dead Violet Dead cell stain kit (dilution: 1:750; Thermo Fisher Scientific). Cells were washed with 2% FBS and phosphate-buffered saline (PBS) and centrifuged at 2000rpm for 5pm. The pellet was resuspended in 100ul 2% FBS and PBS and incubated in Fc block for 5min in that dark at 4C to block nonspecific binding of antibodies. Cells then were incubated for 30 minutes at 4C in the dark with antibodies against cell surface markers. PE-conjugated anti-CD133 (dilution: 1:50; clone: 315-2C11), APC/Fire 700 anti-CD90.2-(dilution: 1:100; clone 5E10), FITC conjugated anti CD45 (dilution: 1:100; clone QA17A26), and APC/Fire 750 conjugated anti E-Cadherin (dilution: 1:100; clone DECMA-1. All antibodies purchase from BioLegend (CA, USA). Samples were washed in 1mL 2% FBS and PBS and centrifuged at 2000rpm for 5mins. Pellets were resuspended in 400-500ul of cold adv DMEM, 10% FBS, and DNase 1, and the single cells were run on a BD LSRFortessa flow cytometer (BD Biosciences). Data were analyzed using FlowJo software (Tree Star, OR).

### RNA-seq Data pre-processing

Reads from the RNA-Sequencing data were aligned to the Mus musculus genome assembly GRCm38 (mm10) using the STAR software^26^. Duplicated aligned reads were marked and removed using the Picardtools software. The gene expression count data were extracted using the HTseq software. The raw count data were normalized, followed by log2 transformation. We filtered out genes with mean read counts < 6. All data preprocessing was performed using R software.

### Differential gene expression analysis

Genes with low counts were removed, and “expressed genes” were defined as those with at least 6 counts in 6 samples. VST was used to normalize RNA-seq across the 12 samples. PCA plots were used as an unbiased approach to visualize the distribution of clusters by age. DESeq2^ref 27^ was performed to address age-related changes in overall levels of gene expression in young vs. aged, and differentially expressed genes were defined as p-value < 0.05 and fold-change cutoff ≥ 1.5.

### ANOVA

One-way ANOVA was calculated in R using car v.3.0.10^ref 28^. Mean square difference between and within groups were calculated. Obtained F values were compared with the critical value in the F table to obtain P values. Inter-group differences were significant (p < 0.05) when the F value exceeded the critical F value for the given degrees of freedom.

### Pathway enrichment

Pathway enrichment analysis for differentially expressed and rhythmic genes was performed with Reactome^29^ using a hypergeometric statistical test and Benjamini-Hochberg FDR correction^30^. Redundant pathway terms were merged to create a parent term.

### Circadian expression analysis

Meta2D cycling algorithm was used to search for genes whose expression cycle in a circadian manner8. Genes were considered cycling if the combined p values were 0.05 or lower for all three algorithms incorporated in Meta2D (ARSER, JTK_CYCLE and Lomb-Scargle).

### Fractional cell abundance

Imputation of fractional abundances of 64 immune and stromal cell types was carried out using immunedeconv package in R. Immunedeconv provides a unified access to 10 most used computational methods for estimating immune cell fractions from bulk RNA sequencing data (quantiseq, cibersort, xcell etc.)^31^.

### Gene panels and Datasets

Fibroblast gene panels were created from ref 32. Fibroblast responsive genes (FRGs) were identified by employing the fibroblast gene panel as input and annotated pathways in Reactome as matching criterion. The analysis was carried out in Cytoscape v.3.4.0. Hub proteins in a network were identified using CytoHubba (MCC, Neighborhood connectivity) plugin in Cytoscape (Ref).

GTeX datasets for human pancreas were obtained from GTeX Analysis V9 (dbGaP accession phs000424.v9). Time stamped human pancreas datasets were obtained from Ref 10 as transcripts per million (TPM).

## Notes

### Competing Interest Statement

The authors have declared no competing interest.

## References

1. Merrow, M., Spoelstra, K., & Roenneberg, T. (2005). The circadian cycle: daily rhythms from behaviour to genes. EMBO reports, 6(10), 930–935. https://doi.org/10.1038/sj.embor.7400541

2. Sanchez, R. E. A., Kalume, F., & de la Iglesia, H. O. (2022). Sleep timing and the circadian clock in mammals: Past, present and the road ahead. Seminars in cell & developmental biology, 126, 3–14. https://doi.org/10.1016/j.semcdb.2021.05.034

3. Mirzaei, K., Xu, M., Qi, Q., de Jonge, L., Bray, G. A., Sacks, F., & Qi, L. (2014). Variants in glucose- and circadian rhythm-related genes affect the response of energy expenditure to weight-loss diets: the POUNDS LOST Trial. The American journal of clinical nutrition, 99(2), 392–399. https://doi.org/10.3945/ajcn.113.072066

4. Oishi, K., Shirai, H., & Ishida, N. (2005). CLOCK is involved in the circadian transactivation of peroxisome-proliferator-activated receptor alpha (PPARalpha) in mice. The Biochemical journal, 386(Pt 3), 575–581. https://doi.org/10.1042/BJ20041150

5. Meijer, J. H., Groos, G. A., & Rusak, B. (1986). Luminance coding in a circadian pacemaker: the suprachiasmatic nucleus of the rat and the hamster. Brain research, 382(1), 109–118. https://doi.org/10.1016/0006-8993(86)90117-4

6. Meijer, J. H., Rusak, B., & Gänshirt, G. (1992). The relation between light-induced discharge in the suprachiasmatic nucleus and phase shifts of hamster circadian rhythms. Brain research, 598(1-2), 257–263. https://doi.org/10.1016/0006-8993(92)90191-b

7. Barclay, J. L., Tsang, A. H., & Oster, H. (2012). Interaction of central and peripheral clocks in physiological regulation. Progress in brain research, 199, 163–181. https://doi.org/10.1016/B978-0-444-59427-3.00030-7

8. Abraham, U., Schlichting, J. K., Kramer, A., & Herzel, H. (2018). Quantitative analysis of circadian single cell oscillations in response to temperature. PloS one, 13(1), e0190004. https://doi.org/10.1371/journal.pone.0190004

9. Rudic, R. D., McNamara, P., Curtis, A. M., Boston, R. C., Panda, S., Hogenesch, J. B., & Fitzgerald, G. A. (2004). BMAL1 and CLOCK, two essential components of the circadian clock, are involved in glucose homeostasis. PLoS biology, 2(11), e377. https://doi.org/10.1371/journal.pbio.0020377

10. Shimba, S., Ishii, N., Ohta, Y., Ohno, T., Watabe, Y., Hayashi, M., Wada, T., Aoyagi, T., & Tezuka, M. (2005). Brain and muscle Arnt-like protein-1 (BMAL1), a component of the molecular clock, regulates adipogenesis. Proceedings of the National Academy of Sciences of the United States of America, 102(34), 12071–12076. https://doi.org/10.1073/pnas.0502383102

11. Marcheva, B., Ramsey, K. M., Buhr, E. D., Kobayashi, Y., Su, H., Ko, C. H., Ivanova, G., Omura, C., Mo, S., Vitaterna, M. H., Lopez, J. P., Philipson, L. H., Bradfield, C. A., Crosby, S. D., JeBailey, L., Wang, X., Takahashi, J. S., & Bass, J. (2010). Disruption of the clock components CLOCK and BMAL1 leads to hypoinsulinaemia and diabetes. Nature, 466(7306), 627–631. https://doi.org/10.1038/nature09253

12. Lee, J., Moulik, M., Fang, Z., Saha, P., Zou, F., Xu, Y., Nelson, D. L., Ma, K., Moore, D. D., & Yechoor, V. K. (2013). Bmal1 and β-cell clock are required for adaptation to circadian disruption, and their loss of function leads to oxidative stress-induced β-cell failure in mice. Molecular and cellular biology, 33(11), 2327–2338. https://doi.org/10.1128/MCB.01421-12

13. Wolff, C. A., Gutierrez-Monreal, M. A., Meng, L., Zhang, X., Douma, L. G., Costello, H. M., Douglas, C. M., Ebrahimi, E., Pham, A., Oliveira, A. C., Fu, C., Nguyen, A., Alava, B. R., Hesketh, S. J., Morris, A. R., Endale, M. M., Crislip, G. R., Cheng, K. Y., Schroder, E. A., Delisle, B. P., Esser, K. A. (2023). Defining the age-dependent and tissue-specific circadian transcriptome in male mice. Cell reports, 42(1), 111982. https://doi.org/10.1016/j.celrep.2022.111982

14. Hood, S., & Amir, S. (2017). The aging clock: circadian rhythms and later life. The Journal of clinical investigation, 127(2), 437–446. https://doi.org/10.1172/JCI90328

15. Davidson, A. J., Sellix, M. T., Daniel, J., Yamazaki, S., Menaker, M., & Block, G. D. (2006). Chronic jet-lag increases mortality in aged mice. Current biology : CB, 16(21), R914–R916. https://doi.org/10.1016/j.cub.2006.09.058

16. Inokawa, H., Umemura, Y., Shimba, A., Kawakami, E., Koike, N., Tsuchiya, Y., Ohashi, M., Minami, Y., Cui, G., Asahi, T., Ono, R., Sasawaki, Y., Konishi, E., Yoo, S. H., Chen, Z., Teramukai, S., Ikuta, K., & Yagita, K. (2020). Chronic circadian misalignment accelerates immune senescence and abbreviates lifespan in mice. Scientific reports, 10(1), 2569. https://doi.org/10.1038/s41598-020-59541-y

17. Welz, P. S., & Benitah, S. A. (2020). Molecular Connections Between Circadian Clocks and Aging. Journal of molecular biology, 432(12), 3661–3679. https://doi.org/10.1016/j.jmb.2019.12.036

18. Birch H. L. (2018). Extracellular Matrix and Ageing. Sub-cellular biochemistry, 90, 169–190. https://doi.org/10.1007/978-981-13-2835-0_7

19. Wu, G., Anafi, R. C., Hughes, M. E., Kornacker, K., & Hogenesch, J. B. (2016). MetaCycle: an integrated R package to evaluate periodicity in large scale data. Bioinformatics (Oxford, England), 32(21), 3351–3353. https://doi.org/10.1093/bioinformatics/btw405

20. Shilts, J., Chen, G., & Hughey, J. J. (2018). Evidence for widespread dysregulation of circadian clock progression in human cancer. PeerJ, 6, e4327. https://doi.org/10.7717/peerj.4327

21. Schüler, S. C., Kirkpatrick, J. M., Schmidt, M., Santinha, D., Koch, P., Di Sanzo, S., Cirri, E., Hemberg, M., Ori, A., & von Maltzahn, J. (2021). Extensive remodeling of the extracellular matrix during aging contributes to age-dependent impairments of muscle stem cell functionality. Cell reports, 35(10), 109223. https://doi.org/10.1016/j.celrep.2021.109223

22. Velez-Delgado, A., Donahue, K. L., Brown, K. L., Du, W., Irizarry-Negron, V., Menjivar, R. E., Lasse Opsahl, E. L., Steele, N. G., The, S., Lazarus, J., Sirihorachai, V. R., Yan, W., Kemp, S. B., Kerk, S. A., Bollampally, M., Yang, S., Scales, M. K., Avritt, F. R., Lima, F., Lyssiotis, C. A., Pasca di Magliano, M. (2022). Extrinsic KRAS Signaling Shapes the Pancreatic Microenvironment Through Fibroblast Reprogramming. Cellular and molecular gastroenterology and hepatology, 13(6), 1673–1699. https://doi.org/10.1016/j.jcmgh.2022.02.016

23. Talamanca, L., Gobet, C., & Naef, F. (2023). Sex-dimorphic and age-dependent organization of 24-hour gene expression rhythms in humans. Science (New York, N.Y.), 379(6631), 478–483. https://doi.org/10.1126/science.add0846

24. Valentinuzzi VS, Scarbrough K, Takahashi JS, Turek FW. (1997). Effects of aging on the circadian rhythm of wheel-running activity in C57BL/6 mice. Am J Physiol. Dec;273(6):R1957–64

25. Possidente, B., S. McEldowney, and A. Pabon. (1995). Aging length-ens circadian period for wheel-running activity in C57BL/6 mice.Physiol. Behav. 57: 575–579

26. Dobin, A., Davis, C. A., Schlesinger, F., Drenkow, J., Zaleski, C., Jha, S., Batut, P., Chaisson, M., & Gingeras, T. R. (2013). STAR: ultrafast universal RNA-seq aligner. Bioinformatics (Oxford, England), 29(1), 15–21. https://doi.org/10.1093/bioinformatics/bts635

27. Love, M. I., Huber, W., & Anders, S. (2014). Moderated estimation of fold change and dispersion for RNA-seq data with DESeq2. Genome biology, 15(12), 550. https://doi.org/10.1186/s13059-014-0550-8

28. Fox J, et al. Sage 2019;3rd edition.

29. Fabregat, A., Sidiropoulos, K., Viteri, G., Forner, O., Marin-Garcia, P., Arnau, V., D’Eustachio, P., Stein, L., & Hermjakob, H. (2017). Reactome pathway analysis: a high-performance in-memory approach. BMC bioinformatics, 18(1), 142. https://doi.org/10.1186/s12859-017-1559-2

30. Jassal, B., Matthews, L., Viteri, G., Gong, C., Lorente, P., Fabregat, A., Sidiropoulos, K., Cook, J., Gillespie, M., Haw, R., Loney, F., May, B., Milacic, M., Rothfels, K., Sevilla, C., Shamovsky, V., Shorser, S., Varusai, T., Weiser, J., Wu, G., D’Eustachio, P. (2020). The reactome pathway knowledgebase. Nucleic acids research, 48(D1), D498–D503. https://doi.org/10.1093/nar/gkz1031

31. Sturm, G., Finotello, F., Petitprez, F., Zhang, J. D., Baumbach, J., Fridman, W. H., List, M., & Aneichyk, T. (2019). Comprehensive evaluation of transcriptome-based cell-type quantification methods for immuno-oncology. Bioinformatics (Oxford, England), 35(14), i436–i445. https://doi.org/10.1093/bioinformatics/btz363

32. Muhl, L., Genové, G., Leptidis, S., Liu, J., He, L., Mocci, G., Sun, Y., Gustafsson, S., Buyandelger, B., Chivukula, I. V., Segerstolpe, Å., Raschperger, E., Hansson, E. M., Björkegren, J. L. M., Peng, X. R., Vanlandewijck, M., Lendahl, U., & Betsholtz, C. (2020). Single-cell analysis uncovers fibroblast heterogeneity and criteria for fibroblast and mural cell identification and discrimination. Nature communications, 11(1), 3953. https://doi.org/10.1038/s41467-020-17740-1

